# Reconstruction of artificial nuclei with nuclear import activity in living mouse oocytes

**DOI:** 10.1101/2024.05.03.592133

**Authors:** Nao Yonezawa, Tomoko Shindo, Haruka Oda, Hiroshi Kimura, Yasushi Hiraoka, Tokuko Haraguchi, Kazuo Yamagata

## Abstract

The cell nucleus is a dynamic structure repeating disassembly and reformation during mitosis. Reformation of the nucleus is essential for cell proliferation, and therefore, the factors required for nuclear reformation are fundamental for eukaryotes. Although various factors have been identified in *in vitro* systems using frog egg extracts and *in vivo* imaging of somatic cells, little is known about the factors required for the formation of functional nuclear structures in living mouse eggs. To identify such factors, we used a reconstruction approach to construct an artificial nucleus around DNA in mouse eggs. T4 phage DNA (166 kbp) was microinjected into living mouse oocytes. Amounts of DNA injected and injection timing were examined to determine the conditions appropriate for the formation of functional nuclei. Microinjection of 100-500 ng/µl DNA during metaphase through telophase of the second meiosis, but not the subsequent interphase, was important for the formation of artificial nucleus. This T4 DNA-derived artificial nucleus had the structure of nuclear lamina and nuclear pore complexes, and nuclear transport activity, similar to natural nuclei. These results suggest that exogenous DNA can form a functional nucleus in mouse oocytes, regardless of the sequence or the source of DNA.

## Introduction

In eukaryotes, genomic DNA forms nucleosomes by binding with histones (two copies of H2A, H2B, H3, H4), and is packed as chromatin in the nucleus. Biological phenomena, such as cell proliferation and development, proceed normally through precise regulation of gene expression. Such precise control of gene expression is achieved in the presence of the nuclear structures, comprising the nuclear envelope (NE), nuclear bodies, and chromatin. Their functions accomplished by nuclear transport through the nuclear pore complex (NPC). The NE comprised of a double membrane structure (inner and outer nuclear membranes), the NPC, and nuclear lamina. The NPC is a large protein complex formed in fenestrated structures between the inner and outer nuclear membranes, and mediates macromolecular transport between the nucleus and cytoplasm. Therefore, nucleocytoplasmic transport activity through the NPC is essential for establishing the nuclear functions, such as regulation of gene expression.

How the functional nucleus is formed is a fundamental question for eukaryotic biology. Studies to identify factors that are required for nuclear formation have been made over the past several decades. One of the successful breakthroughs in biochemical identification of factors was the development of an experimental system using *Xenopus* oocyte extracts (Lohka and Masui, 1983). This system makes it possible to determine the factors required of the NE formation by adding target factor-binding beads to the extract or by adding sperm chromatin to the target factor-depleted extract. Using this system, DNA (Heald et al., 1996), Ran (Zhang & Clarke., 2000), and importin β (Zhang et al., 2002) have been identified as a sole factor to reconstruct the artificial nucleus *in vitro*. More recently, it has been reported that nucleosomes, rather than DNA, are the effective factor for NE assembly (Zierhut et al., 2014). In addition to these *in vitro* experiments, *in vivo* studies using live-cell imaging are also being conducted to explore the factors responsible for NE formation. Barrier-to-autointegration factor (BAF) has been reported to be a factor responsible for NE reformation during the telophase by recruiting LEM domain NE proteins, such as emerin, and Lem2, to the “core region” of the chromosome mass (Haraguchi et al., 2001; Haraguchi et al., 2008), and also for the NE repair during the interphase (Halfmann & Roux, 2021; Kono et al., 2022). Coordination of phosphorylation and dephosphorylation is important for NE reformation during telophase (Vagnarelli et al., 2011; Asencio et al., 2012; Snyers et al., 2018). Endosomal sorting complexes required for the transport-III (ESCRT-III) complex is involved in sealing to the membrane gap of the assembling NE at the end of mitosis (Olmos et al., 2015; Vietri et al., 2015). The interaction of Elys, a nucleoporin, with nucleosomes promotes NPC assembly in tissue-cultured cells (Clever et al., 2012), and also in mouse oocytes (Inoue et al., 2014).

Mouse oocytes provide an excellent experimental system for studying the process of nuclear formation in the living state, enabling to monitor nuclear disassembly and assembly over a short period of time during the process of fertilization and development (Yamagata et al., 2013; Yao et al., 2018; Hatano et al., 2022). Fertilization is the first process in ontogeny, in which two independent cell nuclei, male and female pronuclei are formed (Okada &Yamaguchi, 2017). Although various nuclear functions, such as nuclear transport and transcription, are acquired progressively, little is known about the factors necessary for the formation of functional pronuclear structures in eggs. To identify such factors required for functional pronuclear formation, we used reconstitution approaches in living mouse oocytes by combining *in vitro* approaches. In the previous report, we introduced DNA-coated beads into mouse fertilized-eggs, and found that DNA-coated beads had the ability to form the nucleus with NPCs in mouse eggs (Suzuki et al, 2019). However, this reconstructed nucleus lacked nuclear transport activity. In this paper, to understand factors to form functional NE, we attempt to reconstruct the artificial nucleus with nuclear transport activity in living mouse oocytes.

## Results

### 166 kbp-long T4 DNA forms a single nucleus in living mouse oocytes

We first investigated the preferred DNA length for the formation of artificial nuclei (Figure 1). T4 phage genomic DNA (T4 DNA; approximately 166 kbp in length) and plasmid pGADT7 DNA (pGADT7 DNA; ∼8 kbp in length) were tested as a DNA source. Unfertilized oocytes arrested at metaphase II (MII oocytes) were microinjected with mRNA encoding histone H2B-mCherry as a fluorescent probe to visualize chromatin (Figure 1a). The oocytes were activated with strontium to mimic fertilization, and 30∼40 minutes later, the oocytes were microinjected with DNA (approximately 5 picoliters (pl) of DNA solution to each oocyte) (Figure 1a). Just after DNA microinjection, the injected region was clearly seen in the oocyte with T4 DNA under a bright-field microscopy, whereas it was not clear in those with pGADT7 DNA (Figure 1b and movie S1). The microinjected region was also clearly seen with λDNA (approximately 55 kbp in length; Figure S1a). Sixty minutes after microinjection, the injected sites were still clearly visible in T4 DNA-injected oocytes under the bright-field microscopy, and the microinjected DNA was also detected with Hoechst 33342 as one large mass in the injected region (right panels in Figure 1c). Similar results were obtained with λDNA (Figure S1a). In contrast, in pGADT7 DNA-injected oocytes, the injected site was not clearly visible by the bright-field microscopy and the location of pGADT7 DNA was not detected with Hoechst 33342 (left panels in Figure 1c). To further clarify the fate of the injected DNA, live-cell imaging was performed for oocytes fluorescently-labeled with histone H2B-mCherry. The results showed that in oocytes injected with pGATD7 DNA, histone H2B-mCherry signal was hardly detected until 60 min after injection, multiple small dots with H2B-mCherry signals appeared at 90 min after injection, and then the number of dots increases until 300 min (Figure 1d). In contrast, in oocytes injected with T4 DNA, a single mass of H2B-mCherry signals was detected in the injection site 30 min after injection (arrowheads, Figure 1d). The fluorescent mass became larger over time and then formed one large spherical structure with H2B-mCherry signals at 300 min (arrowheads, Figure 1d). Within the structure, the distribution of H2B-mCherry was not uniform, and punctate accumulations of H2B-mCherry signals were observed, as was seen in maternal pronuclear formation (“T4DNA”, arrowheads at 300 min in Figure 1d). Based on this localization pattern of H2B-mCherry, this large spherical structure was morphologically similar to the maternal pronucleus, suggesting that the injected T4 DNA formed nucleus-like structures. Hereafter, we called this structure as the artificial nucleus. Similarly, injection of λDNA resulted in the formation of the artificial nucleus with H2B-mCherry signals (Figure S1a). These results suggest that the length of DNA may be one of the factors influencing the construction of artificial nuclei within cells. Based on this, we decided to use T4 DNA for the following experiments.

**Figure 1.**
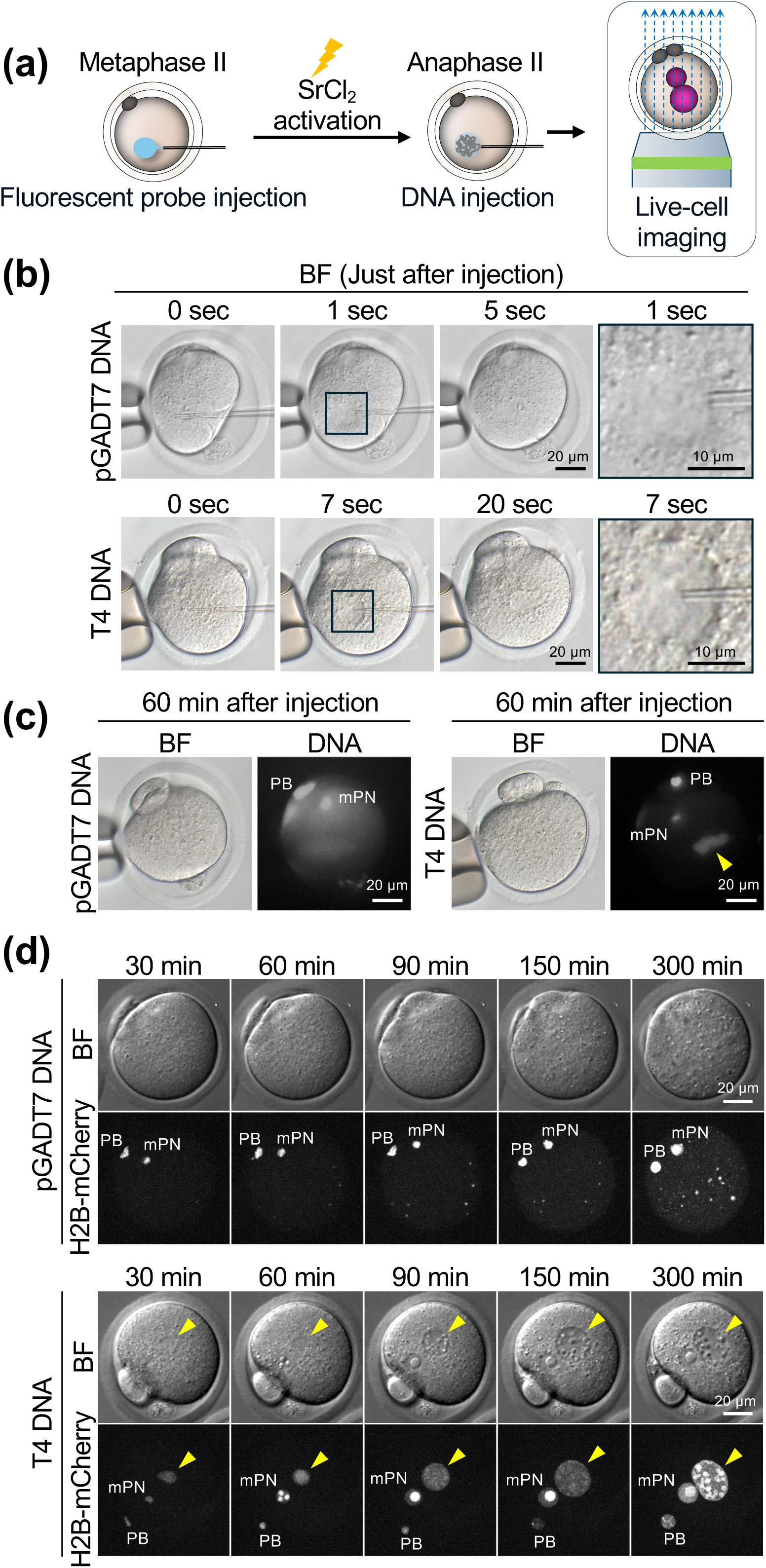
Microinjection of DNA into strontium activated mouse oocytes. (a) Schematic illustration of DNA microinjection. Unfertilized mouse oocytes arrested at metaphase II were microinjected with mRNA encoding fluorescent protein of interest as a fluorescent probe. The oocytes were activated with SrCl_2_ to mimic fertilization, injected with DNA solution, and subjected to live-cell imaging using fluorescence microscopy. (b) Representative time-lapse bright-field (BF) microscopic images of living mouse oocytes during microinjection of pGADT7 DNA (upper panels) and T4 DNA (lower panels). The numbers above each image indicate the time in seconds after DNA injection. The rightmost panel is an enlarged view of the black frame on the left. Scale bars, 20 µm (left panels) and 10 µm (enlarged right panel). (c) Representative images of living mouse oocytes 60 min after microinjection of pGADT7 DNA (left panels) and T4 DNA (right panels). The left side of each image set is a BF image, and the right side is a fluorescent image. DNA was stained with 0.5 µg/ml Hoechst 33342 for 30 min before image acquisition. The concentration of injected DNA is 500 ng/µl. Arrowheads indicate injected T4 DNA. PB, polar body. mPN, maternal pronucleus. Scale bars, 20 µm. (d) Time-lapse images of living mouse oocytes injected with pGADT7 DNA (upper panels) and T4 DNA (lower panels). BF and H2B-mCherry (maximum projection images) are shown. The numbers above each image indicate the time in minutes after DNA injection. Arrowheads indicate the region of injected T4 DNA. PB, polar body. mPN, maternal pronucleus. Scale bars, 20 µm.

### DNA injection timing is important for the formation of artificial nucleus

Next, we examined the preferred timing of the DNA injection for nuclear formation during the second meiotic cell cycle (Figure 2). To visualize nuclear formation, unfertilized oocytes were injected with mRNA encoding H2B-mCherry as a probe, and then injected with T4 DNA at various timings before or after strontium activation (30 min before, 30, 120, and 360 min after activation) (Figure 2a). The corresponding cell cycle stages of the DNA injection were second meiotic metaphase (metaphase II), second meiotic anaphase (anaphase II), second meiotic telophase (telophase II) and interphase, respectively (Figure 2a). The oocytes were observed by fluorescence microscopy in the living state at 300 min after injection (Figure 2b). When DNA was injected into metaphase II oocytes, one artificial nucleus with H2B-mCherry signals formed around the injected DNA (arrowheads in Metaphase II panels, Figure 2b). In the artificial nucleus, punctate accumulations of H2B-mCherry were observed as found in the natural maternal pronucleus (H2B-mCherry panels, Figure 2b). However, under this experimental condition, the DNA injection caused significant errors in the polar body extrusion in almost all oocytes (green dashed region, Figure 2b). The size of the released polar bodies varied, and some had the morphology of two-cell stage embryos (compare 2^nd^ PB in Metaphase II with those in Anaphase II and Telophase II, Figure 2b). In contrast, when DNA was injected at anaphase II and telophase II, artificial nuclei with H2B-mCherry signals were formed around the DNA and no obvious polar body abnormalities were observed (arrowheads in Anaphase and Telophase panels, Figure 2b). In contrast, however, in oocytes injected at the interphase, no H2B-mCherry signals appeared in the spherical-shaped DNA aggregates with Hoechst signals formed at the injected site (Interphase, Figure 2b). Quantitative analysis of H2B-mCherry signals in artificial nuclei revealed that recruitment of histone H2B to DNA was significant in injection at metaphase II, anaphase II and telophase II, but not at interphase (Figure 2c). These results suggest that the timing of DNA injection is important for the formation of artificial nuclei in living mouse oocytes and that DNA must be introduced during metaphase II through telophase II. Based on these results, we used anaphase II oocytes for further experiments unless otherwise stated.

**Figure 2.**
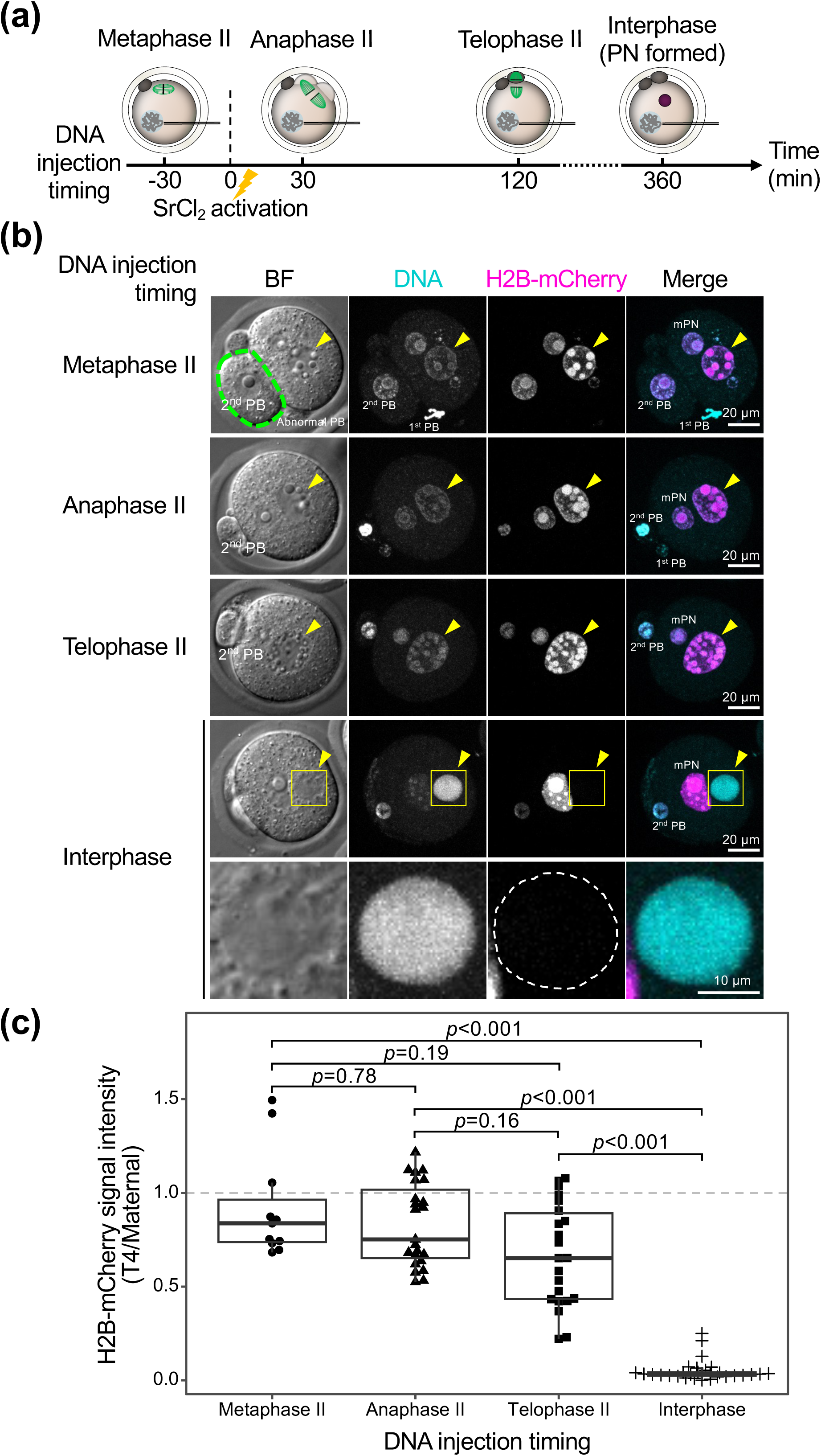
Effects of timing of DNA microinjection on the nuclear formation. (a) Schematic diagram of DNA microinjection timing. DNA was injected into living mouse oocytes at the indicated timings with SrCl_2_ activation set at 0 min. DNA microinjection was performed at metaphase, anaphase, telophase of meiosis II or interphase (pronucleus (PN) formation stage).(b) Representative microscopic images of BF, DNA and H2B-mCherry in living mouse oocytes 300 min after injecting T4 DNA at metaphase II, anaphase II, telophase II, and interphase. DNA was stained with 0.5 µg/ml Hoechst33342. Single focus images are shown for BF. Maximum projection images for DNA and H2B-mCherry. Yellow arrowheads indicate the region of injected T4 DNA. The bottom panels are magnified views of the yellow squares in the panels immediately above their respective panels. The region surrounded by the green dashed line indicates the position of the second polar body (2^nd^ PB). 1^st^ PB, first polar body. mPN, maternal pronucleus. Scale bars, 20 µm (overview), 10 µm (magnified view). (c) Quantitative analysis of H2B-mCherry signals of T4 DNA artificial nuclei in oocytes shown in (b). The Y axis shows the intensity value of the artificial nucleus relative to the maternal nucleus in arbitrary units. Box-and-whisker plots are shown: the box indicates the median and the upper and lower quartiles; the whisker indicates the range. The number of samples tested, n = 11, 23, 22, and 30 from the left. Significant differences are presented as *p* < 0.05 (Steel-Dwass test).

### DNA concentrations required for the formation of artificial nucleus

We examined the DNA concentrations to be injected (Figure 3). Anaphase II oocytes expressing H2B-mCherry were injected with various concentrations (20, 100, and 500 ng/µl) of DNA, and subjected to the live-cell imaging. Images at 300 min after injection were shown in Figure 3. When oocytes were injected with T4 DNA at a concentration of 20 ng/µl, no structures with Hoechst and H2B-mCherry signals were detected except for those of maternal pronucleus and polar body (Figure 3). In contrast, at higher concentrations (100 and 500 ng/µl), a single spherical structure, as large or slightly larger than the maternal pronucleus, was formed with Hoechst and H2B-mCherry signals (Figure 3). Within the structure, H2B-mCherry was unevenly distributed, similar to its distribution in the maternal pronucleus. These results suggest that a certain level of concentration, i.e., 100 ng/µl or higher, is required for the formation of artificial nuclei.

**Figure 3.**
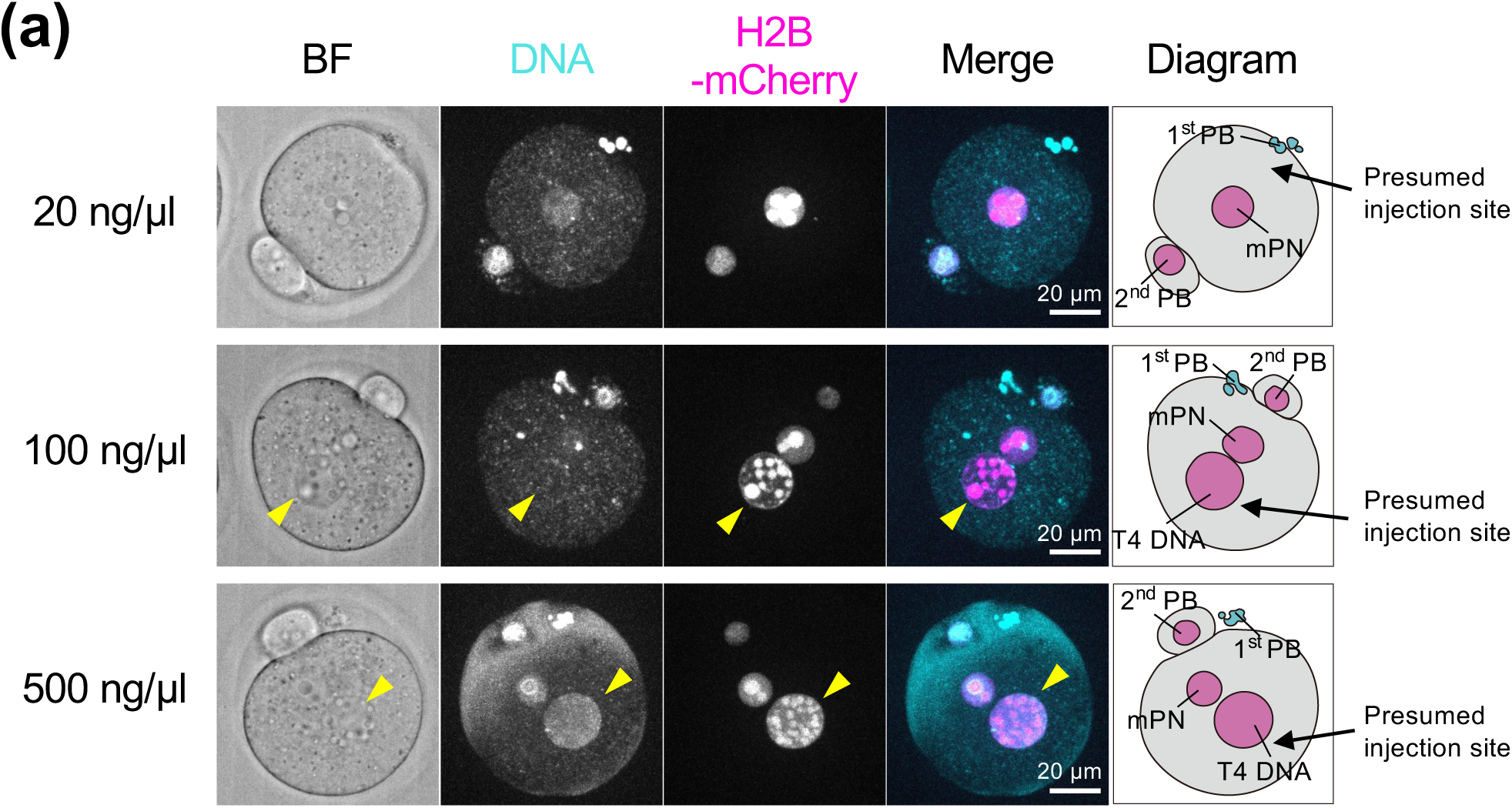
Effects of the DNA amount on the nuclear formation. Representative microscopic images of BF, DNA, H2B-mCherry, and their merge in mouse oocytes. DNA was stained with Hoechst 33342. Images were taken 300 min after injecting anaphase II mouse oocytes with the indicated concentrations of T4 DNA. Single focus images are shown for BF. Maximum projection images for DNA and H2B-mCherry. Rightmost diagrams show positions of polar bodies (PB), maternal pronucleus (mPN), and T4 DNA-injected region (T4 DNA). Scale bar, 20 µm.

### Nucleosome formation in the T4 DNA artificial nucleus

We assessed the integrity of the artificial nucleus constituted by the T4 DNA (Figures 4-6). First, nucleosome formation on the DNA was analyzed (Figure 4). Anaphase II oocytes were injected with 250 ng/µl T4 DNA and incubated for 300 min. After fixation, the oocytes were subjected to immunofluorescent staining analysis. Results showed that the T4 DNA-dependent artificial nuclei were stained with specific antibodies against any of the canonical histones tested (H2A, H3, and H4) with a staining intensity comparable to that of maternal nuclei (Figure 4a-c). All histones H2A, H3, and H4 were ununiformly distributed within the artificial nucleus, similar to the maternal nucleus. Notably, histone-free spherical regions were formed within the artificial nuclei as well as in the maternal nuclei, and accumulation of histones H2A and H3 was observed at the edges of the spherical region. This histone-free region was stained with B23, a component of the nucleolus, by immunostaining with anti-B23 antibody (Figure S2a), suggesting that this spherical histone-free region is likely a structure similar to nucleolar precursor body.

**Figure 4.**
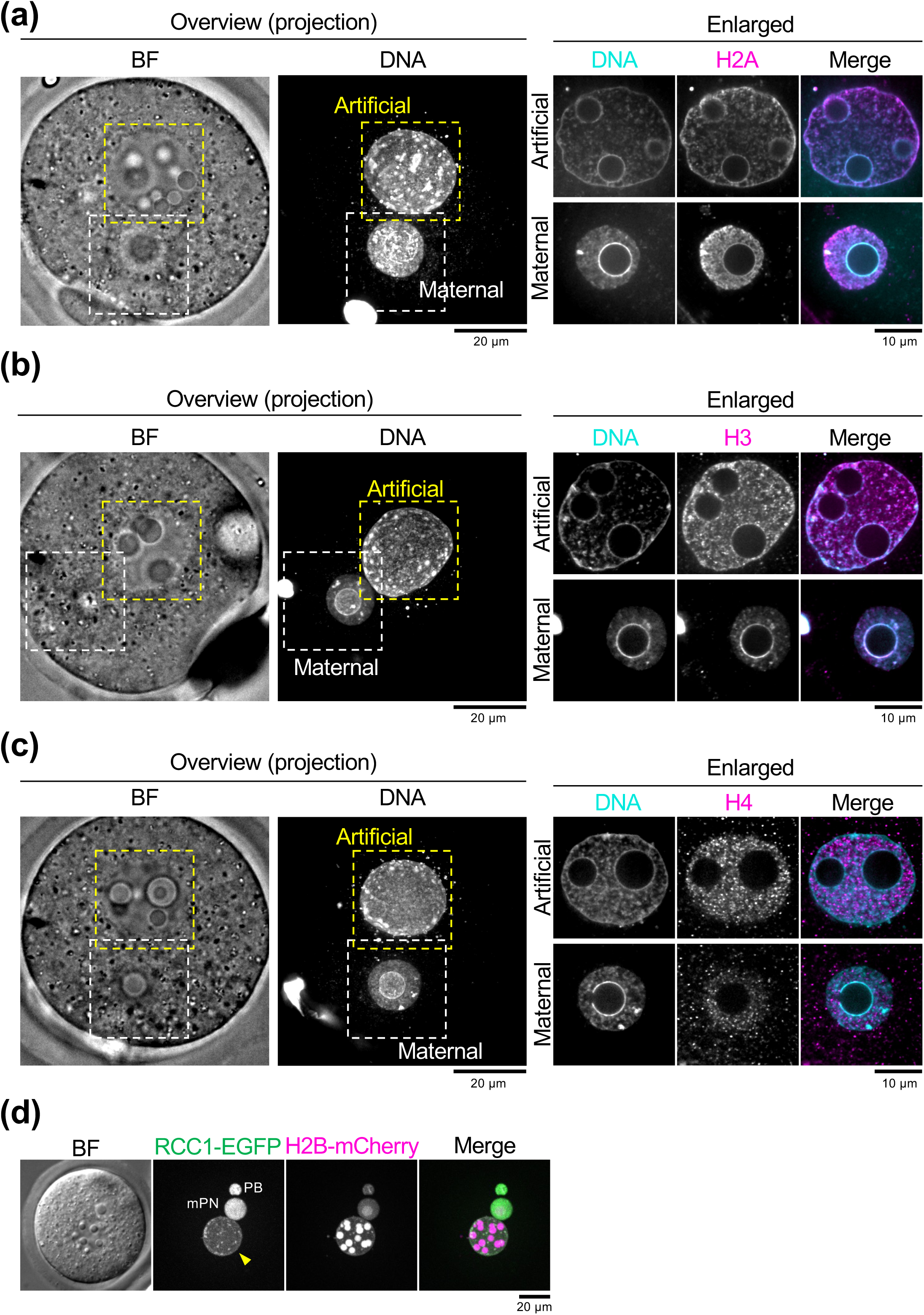
Nucleosome formation on the DNA in the artificial nuclei. (a) Immunostaining images of mouse oocytes using anti-histone H2A antibody. Oocytes were fixed at 300 min after injection of 250 ng/µl T4 DNA at anaphase II. DNA was stained with DAPI. The right panels are enlarged views of the yellow and white frames on the left. Left panels: a BF image is a single focus image and a DNA image is a maximum projection image of 46 images in Z-axis direction. Right panels: single plane images of equatorial plane of the nucleus indicated in the left panels. The upper panels show an artificial nucleus formed by T4 DNA, and the lower panels a maternal pronucleus. The merged image, DNA (cyan) and histones (magenta). Scale bar, 10 µm. (b) Immunostaining images using anti-histone H3 antibody. Same as (a), except that anti-H3 antibody was used. (c) Immunostaining images using anti-histone H4 antibody. Same as (a), except that anti-H4 antibody was used. (d) Microscopic images of living mouse oocytes. Anaphase II oocytes probed with RCC1-EGFP and H2B-mCherry were injected with 250 ng/µl T4 DNA. BF, RCC1-EGFP, and H2B-mCherry images were taken 300 min after DNA injection. Single focus images are shown for BF. Maximum projection images for DNA and H2B-mCherry. The merged image, RCC1-EGFP (green) and H2B-mCherry (magenta). Yellow arrowheads indicate the region of injected T4 DNA. PB, polar body. mPN, maternal pronucleus. Scale bar, 20 µm.

**Figure 5.**
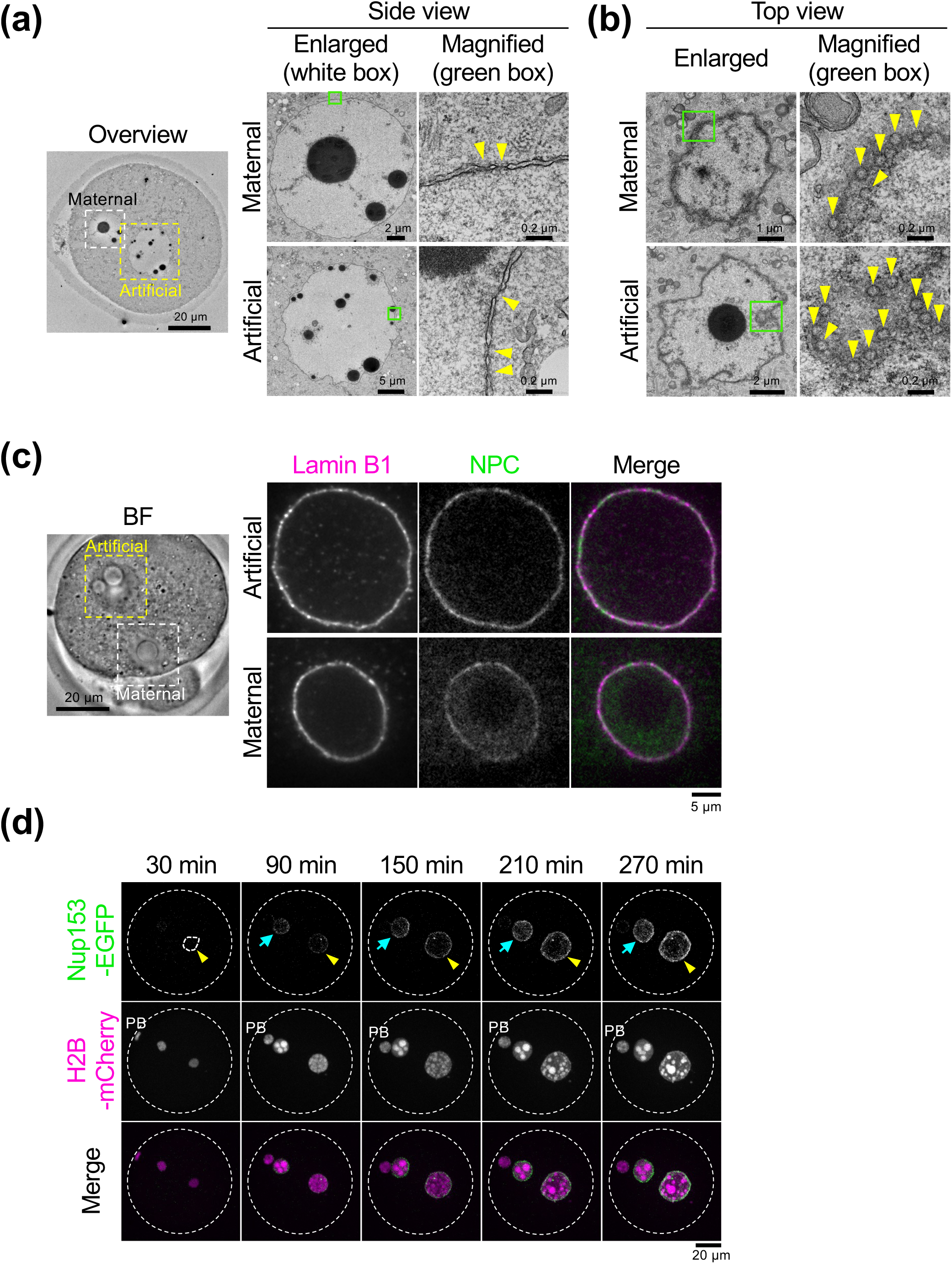
Nuclear envelope structures of the artificial nuclei. (a) Electron microscopy (EM) images of mouse oocytes 300 min after injecting 250 ng/µl T4 DNA at anaphase II. The leftmost image is an overview EM image of equatorial section of the nuclei. Enlarged images show the images of the yellow and white frames in the overview. Magnified images show images of the green-boxed region in the enlarged panel. Upper and lower images are maternal pronuclei and T4 DNA artificial nuclei, respectively. Yellow arrowheads indicate positions of the nuclear pore complex (NPC)-like structures. Scale bars are indicated in each panel. (b) Different section images of the same sample as in (a). Surface section of the maternal nucleus and artificial nucleus. Yellow arrowheads indicate positions of the NPC-like structures. Scale bars are indicated in each panel. (c) Immunostaining images of mouse oocytes using anti-lamin B1 and anti-NPC (mAb414) antibodies. Oocytes were fixed at 300 min after injecting 250 ng/µl T4 DNA into anaphase II oocytes. The leftmost image is a BF image. The right upper panels show fluorescence images of artificial nuclei (yellow-boxed region) and the lower panels maternal nuclei (white-boxed region). The merged image, lamin B1 (magenta) and NPC (green). The images are single planes of the nuclei. Scale bar, 5 µm. (d) Time-lapse fluorescence images of living mouse oocytes. Anaphase II oocytes expressing Nup153-EGFP and H2B-mCherry were injected with 250 ng/µl T4 DNA. The numbers above each image indicate the time in minutes after DNA injection. Maximum projection images are shown. Yellow arrowheads and blue arrows indicate the regions of injected T4 DNA and maternal pronucleus, respectively. The merged image, Nup153-EGFP (green) and H2B-mCherry (magenta). PB, polar body. Scale bars, 20 µm.

**Figure 6.**
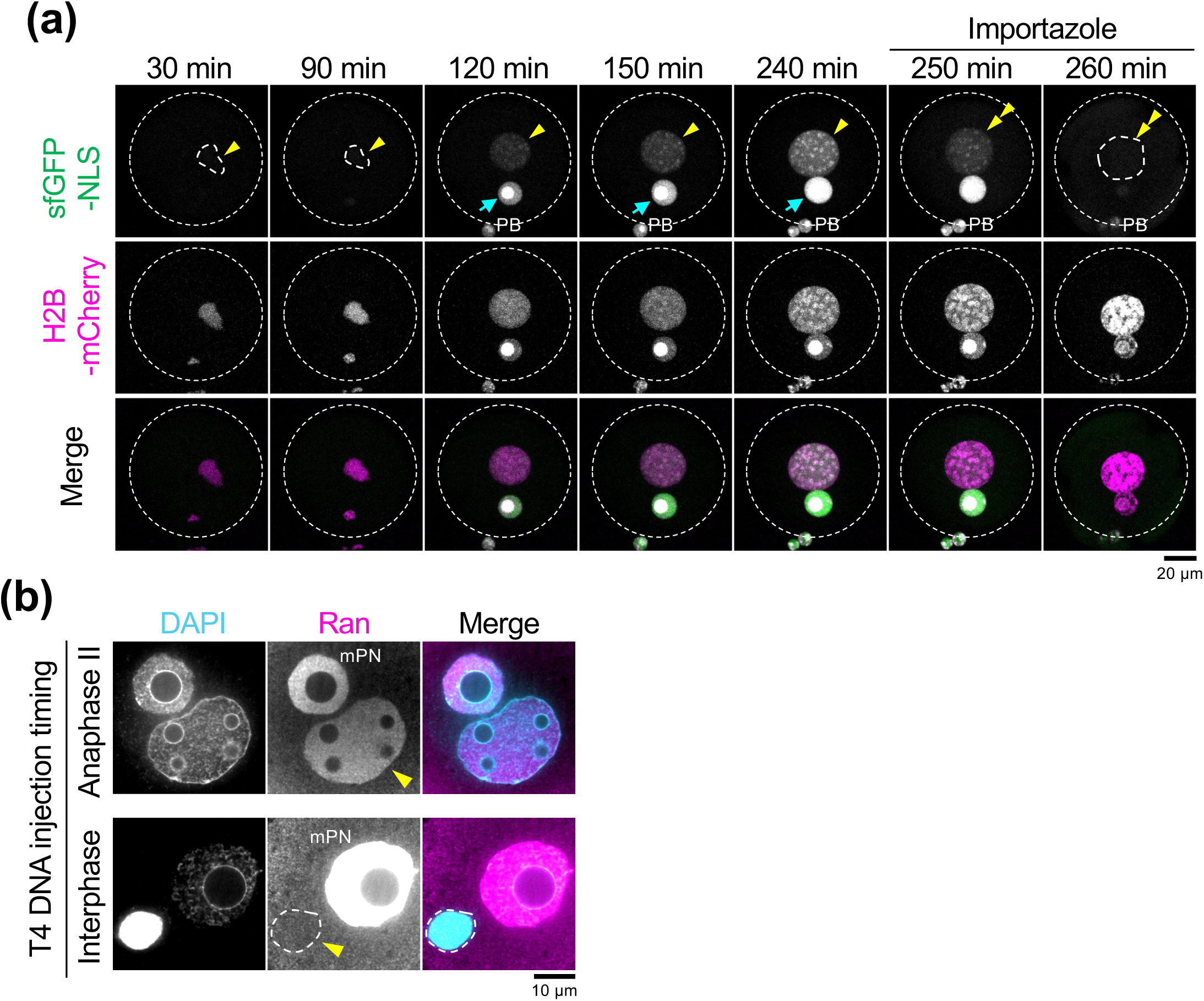
Nuclear transport activity of the artificial nuclei. (a) Time-lapse fluorescence images of living mouse oocytes. Anaphase II oocytes expressing sfGFP-NLS and H2B-mCherry were injected with 250 ng/µl T4 DNA. The numbers above each image indicate the time in minutes after DNA injection. Maximum projection images are shown. The merged image, sfGFP-NLS (green) and H2B-mCherry (magenta). Yellow arrowheads and blue arrows indicate the regions of injected T4 DNA and maternal pronucleus, respectively. Importazole, an inhibitor of importin β, was added to oocytes at 250 min. Double arrowheads indicate the T4 DNA region after importazole addition. Scale bars, 20 µm. (b) Immunostaining images of mouse oocytes using anti-Ran antibody. Images were taken 300 min after injecting 250 ng/µl T4 DNA into anaphase II oocytes (upper panels) or interphase oocytes (lower panels). DNA was stained with DAPI. The merged image, DNA (cyan) and Ran (magenta). Yellow arrowheads indicate the region of injected T4 DNA. The region surrounded by the white dashed line indicates the position of the injected DNA. The images are single planes of the nuclei. Scale bars, 10 µm.

RCC1 is known to bind to acidic patches in nucleosome structures (Makde et al., 2010). Therefore, to confirm whether T4 DNA forms nucleosomes, we examined the presence of RCC1. Anaphase II oocytes expressing RCC1-EGFP and H2B-mCherry were injected with 250 ng/µl DNA, and subjected to the live-cell imaging. Results showed that RCC1 was positive in the artificial nucleus, although the RCC1-EGFP signal was relatively lower than that of the maternal nucleus (Figure 4d). This result suggests that T4 DNA forms nucleosomes in the artificial nucleus.

### The NE structure in the T4 DNA artificial nucleus

To further assess the structure of the artificial nuclei formed by T4 DNA, we examined the membrane structure surrounding the artificial nucleus by electron microscopy (Figure 5a, b). Side views of the artificial nucleus showed the structure characteristic of the NE, i.e. a double membrane structure and a constriction structure in between (Figure 5a). This membrane structure was similar to that seen in the maternal pronucleus. In addition, top views showed that there are many pore structures, which were indistinguishable from those in the maternal pronucleus, on the double membrane structures (arrowheads, Figure 5b), suggesting that NPC-like structures were formed on the NE of the artificial nucleus. It is interesting to point out that in addition to these membrane structures, several to ten electron-dense spherical structures were formed in the artificial nucleus (Figure 5a, b). Since electron-dense structures also exist in the maternal nucleus, it may be a structure similar to the nucleolus precursor body. In contrast, the membrane structures around T4 DNA injected at interphase were similar to the endoplasmic reticulum with complex branched and tangled membrane structures, but with no NPC-like structure on the membrane (Figure S2a).

Immunostaining was performed using the specific antibodies, anti-laminB1 and anti-NPC (mAb414) antibodies to the nuclear lamina and NPCs, respectively. Results showed that positive signals of lamin B1 and NPC were observed at the periphery of the T4 DNA artificial nucleus, similar to those observed in the maternal nucleus (Figure 5c). Similar results were obtained with λDNA artificial nuclei (Figure S1b). Furthermore, lamin B1 and NPC signals were located at mutually exclusive positions on the NE in both the artificial and natural maternal pronuclei (Figure 5c). This result is consistent with previous results showing that the nuclear lamina and NPC exist in exclusive locations (Fišerová et al., 2017), indicating that this artificial nucleus has the NE structure indistinguishable from that of the natural nucleus.

We also performed live-cell imaging to understand the process of NE assembly in the artificial nuclei (Figure 5d). Nup153, a component of the NPC, was tagged with EGFP (Nup153-EGFP) and expressed as a probe for the NPCs, along with the H2B-mCherry probe for chromosomes. Anaphase II oocytes were injected with 250 ng/µl T4 DNA, and monitored in a living state by fluorescence microscopy. Results showed that Nup153-EGFP was recruited around the artificial nucleus at the same timing with the maternal nucleus (Figure 5d). These data suggest that T4 DNA has the ability to form the normal structure of the NE. In contrast, Nup153-EGFP signals did not appear around T4 DNA aggregates when the DNA was injected at interphase (Figure S2c).

### T4 DNA artificial nucleus has nuclear transport activity

We evaluated the nuclear import activity that had not been observed in artificial nuclei constructed by DNA beads (Suzuki et al., 2019). To monitor the nuclear transport activity of artificial nuclei, we used superfolder GFP-nucleoplasmin (NPM)-NLS (Oda et al., 2023) (hereafter, designated sfGFP-NLS). sfGFP-NLS mRNA was microinjected into unfertilized oocytes together with the H2B-mCherry mRNA. Fluorescence live-cell imaging showed that H2B-mCherry signals appeared simultaneously in the artificial and maternal nuclei at 30 min after DNA injection (arrowheads, “30 min” panels, Figure 6a). sfGFP-NLS signals appeared in the both nuclei at 120 min, although the signal was weaker in the artificial nucleus than the maternal pronuclei (compare arrowheads with arrows, “120 min” panels, Figure 6a). Furthermore, as time progressed, the sfGFP-NLS signal in artificial nucleus increased, similar to in the natural maternal pronucleus (mPN) (compare arrowheads with arrows, 120-240 min panels, Figure 6a). This result suggests that the artificial nucleus has nuclear transport activity. To confirm whether the increase in sfGFP-NLS signal was caused by the function of importin β, we examined the effect of importazole, an inhibitor of importin β (Soderholm et al., 2011).

Importazole was added to the oocytes immediately after the image acquisition of 240 min in Figure 6. The nuclear accumulation of sfGFP-NLS was inhibited by addition of importazole (double arrowheads in 250 min and 260 min panels, Figure 6a), suggesting that sfGFP-NLS is actively transported to the nucleus by the function of nuclear transport machineries. Notably, sfGFP-NLS signals decreased, suggesting that nuclear export also occurred in the artificial nuclei.

To confirm nuclear transport activity, we also examined the presence of Ran (Ras-related nuclear protein) in the artificial nucleus: Ran is a small GTPase protein essential for the active transport through the NPC. Anaphase II oocytes were injected with T4 DNA to form artificial nuclei. These artificial nuclei were examined by immunostaining analysis using anti-Ran antibody. The results revealed that the artificial nuclei had similar amounts of Ran as the natural maternal pronucleus (“Anaphase II”, Figure 6b). In contrast, no signals of Ran were observed in DNA aggregates when T4 DNA was introduced into interphase oocytes (“Interphase”, Figure 6b). These results suggest that the T4 DNA artificial nucleus actively transport NLS-bearing proteins through the NPC.

## Discussions

We succeeded in constructing the artificial nucleus in mouse oocyte by injection of DNA solution containing long-strand DNA, T4 DNA (∼166 kbp). However, the oocyte was not always capable of forming an artificial nucleus, and some strict conditions were required to produce a functional artificial nucleus. First, the length of DNA was important. Relatively long DNAs (approximately 166 kbp and 54 kbp) formed a single large-sized nucleus-like structure in oocytes, whereas shorter (approximately 8.8 kbp) DNA only formed many scattered puncta (Figures 1d and S1). This difference is probably due to differences in physicochemical properties of DNA depending on its length. The diffusion coefficient is larger for long polymers compared to short polymers (de Gennes, 1979). Furthermore, the rate of DNA diffusion in solution and within the HeLa cell cytoplasm is dependent on DNA size (Lukacs et al., 2000). Therefore, longer DNA tends to remain at the injected site within the cytoplasm without scattering. In addition, cytoplasmic factors, such as membranes, and physical environment of the cytoplasm, such as density and viscosity, may also be involved in the DNA diffusion depending on the length. Moreover, the effect of DNA length on nuclear formation may be explained by phase separation. Recently, it has been reported that relatively longer nucleosome arrays (approximately 12.5 kbp) and nucleosome-wrapped λDNA (approximately 48.5 kbp) tend to form gel- or solid-like aggregates in buffers, whereas shorter nucleosome arrays (approximately 3 kbp) form droplets with high fluidity (Chen et al., 2022). The speed of nucleosome assembly on λDNA in *Xenopus* oocyte extracts is a few seconds (Ladoux et al., 2000). Thus, DNA dispersion depends on the amounts of nucleosomes on the DNA. Our results suggest that slow dispersion of DNA in the ooplasm after injection is important for nuclear formation.

Second, the appropriate DNA concentration to form obvious artificial nuclei was 100– 500 ng/*μ*l (Figure 3). We estimated this amount of DNA compared to the amount of DNA in the natural nucleus as follows. Our microinjection technique uses a quantitative microinjection method with a piezo drive attached to a screw-controlled syringe (Kline, 2009). The volume of solution injected varies in the range of 0.8–1.2 times in each injection (Yamagata, 2010). The volume of the DNA solution injected was estimated to be approximately several picoliters (pl). Therefore, the amount of injected DNA was estimated to be approximately 500 ng/µl x 5 pl = 2.5 pg. The DNA content in a single sperm of mouse is estimated as 2.75 pg as the haploid genome of a mouse is approximately 2.5 billion base pairs. This DNA content is comparable to the measured value 3.36 ± 0.09 pg using particle-induced X-ray emission-based mass spectrometry (Bench et al., 1996). Therefore, the amount of DNA injected in this study is approximately equivalent to the amount of DNA contained in a sperm. This suggests that amounts of DNA comparable to mouse sperm may be suitable for reconstructing the nuclear structure in mouse oocytes.

Third, the timing of DNA injection appears to be critical for nuclear formation. The formation of the nuclear envelope occurs during the telophase. Therefore, passing through the telophase is considered to be important for the formation of the functional nuclear envelope and thus the functional nucleus. Consistent with this hypothesis, our results show that DNA injected during anaphase II (thus, passing through the entire period of telophase) formed functional nuclei with nuclear transport activity (Figures 2 and 6). This hypothesis is also supported by the result that the degree of nuclear formation was relatively high in injection at metaphase II and anaphase II, slightly lower at telophase II, and none in interphase (Figure 2). This result is consistent with previous findings that that λDNA injected into metaphase *Xenopus* oocytes formed functional nuclei with nuclear transport activity (Forbes et al., 1983), whereas DNA-conjugated beads injected into interphase mouse fertilized-egg did not (Suzuki et al., 2019). However, abnormal polar body extrusion was observed in almost all oocytes injected with DNA at metaphase II. This phenomenon may be related to the RanGTP gradient. The RanGTP gradient is formed around the chromosomes by the function of chromatin-bound RCC1, Ran-specific guanine nucleotide exchange factor, and known to be involved in spindle formation in mitosis (reviewed in Kalab & Heald, JCS 2008). Besides spindle formation, RanGTP gradients are involved in many metaphase-specific events during fertilization such as polar body extrusion (Kaláb et al., 2011; Dumont & Verlhac, 2013), the cytoskeletal arrangement during polar body extrusion (Maro & Verlhac, 2002; Dumont & Verlhac, 2013), and sperm DNA localization during sperm-egg fusion (Mori et al., 2021). DNA injection at metaphase II may have induced a perturbation of the RanGTP gradient within metaphase II oocytes, leading to aberrant polar body protrusion. Therefore, DNA injection at anaphase II is appropriate for the artificial nuclear formation in mouse oocytes.

One of the key factors required for the functional nucleus is the nucleosome because the presence of nucleosomes on chromatin is essential for the formation of the functional nuclear envelope in mouse fertilized-eggs (Inoue et al., 2014) and *Xenopus* egg extracts (Zierhut et al., 2014). This notion is supported by our results that T4 DNA artificial nuclei contain nucleosomes comparable to the amount of natural maternal pronuclei (Figure 4). However, despite the presence of nucleosomes and nucleosome-binding protein RCC1, DNA-conjugated beads injected into interphase mouse fertilized-eggs failed to form functional nuclei with nuclear transport activity (Suzuki et al., 2019). This suggests that the presence of nucleosomes and RCC1 is essential, but not sufficient for the formation of functional nuclei. In addition to the presence of these factors, the presence of RanGTP is also absolutely required for the formation of functional nuclei with nuclear transport activity. This is because the RanGTP gradient plays an essential role in nuclear transport (Yoneda, 1997; Yoneda et al., 1999; Izaurralde et al., EMBO J 1997; Görlich et al., 1999), and in the nuclear envelope assembly in *in vitro* experiments using *Xenopus* egg extracts (Hetzer et al., 2000; Walther et al., 2023; Forbes et al., 2015). Consistently, Ran existed in the artificial nucleus bearing nuclear transport activity, but not in the DNA aggregates (Figure 6b) nor the nucleus-like structures induced by DNA-beads (Suzuki et al., 2019). Given that passage through telophase is required for acquirement of nuclear transport activity, events from metaphase to telophase may be necessary for the establishment of the RanGTP gradient in the interphase nuclei. However, such mechanisms remain unclear and future research is necessary. Taken together, the timing of DNA introduction, the amount of DNA, and the physicochemical environment around the DNA are considered to be important for forming the functional nucleus with nuclear transport activity. This study demonstrates that DNA can form a functional nucleus in mouse oocytes, regardless of the sequence or the source of DNA.

## Experimental procedures

### Materials

Importazole was purchased from MedChemExpress (HY-101091; NJ, USA). The reagent was dissolved in DMSO at the concentration of 100 mM and stored at −30°C as a stock solution. To inhibit nuclear transport, oocytes were treated with 100 µM importazole or DMSO as a control.

The pGADT7 vector (pGADT7 AD, 7,987 bp) and T4 DNA (T4GT7, 166,644 bp) was purchased from Clontech (630442; Clontech Laboratories, Inc., Mountain View, CA, USA) and Nippon Gene (318-03971; Tokyo, Japan), respectively.

### Antibodies

Primary antibodies were used as follows: Mouse monoclonal antibodies against histone H2A (D210-3; Medical & Biological Laboratories Co., Ltd., MBL, Nagoya, Japan); histone H3 (MABI0301; Monoclonal Antibody Laboratory Inc., Nagano, Japan); histone H4 (MABI0400; Monoclonal Antibody Laboratory Inc., Nagano, Japan); anti-NPC (mAb414, 902901; BioLegend, CA, USA); anti-Ran (BD610341; BD Biosciences, NJ, USA); anti-B23 (FC82291; Sigma-Aldrich, St. Louis, USA), and a rabbit polyclonal antibody against lamin B1 (ab16048; abcam, Cambridge, UK). Secondary antibodies were used as follows: CF 488-conjugated goat anti-mouse IgG (H+L) antibody (20302-1; Biotium, CA, USA), CF 555-conjugated goat anti-mouse IgG (H+L) antibody (20231-1; Biotium, CA, USA), CF 488-conjugated goat anti-rabbit IgG (H+L) antibody (20019-1; Biotium, CA, USA), CF 555-conjugated goat anti-rabbit IgG (H+L) antibody (20232-1; Biotium, CA, USA).

### Animals

This study was conducted in compliance with the requirements of the Guide for the Care and Use of Laboratory Animals. All animal experiments were approved by the Animal Care and Use Committee at Kindai University (permit number: KABT-2022-012). ICR strain mice (11–18 weeks old) were obtained from Japan SLC, Inc. (Shizuoka, Japan). Room conditions were standardized with temperature maintained at 23°C, relative humidity at 50% and a 12/12 h light/dark cycle. Animals had free access to water and commercial food pellets.

### Oocyte collection and strontium activation

ICR strain female mice (11-18 weeks old) were treated with 10 IU of pregnant mare serum gonadotropin (PMSG, ASKA Animal Health Co., Ltd., Tokyo, Japan) and 10 IU of human chorionic gonadotropin (hCG, ASKA Animal Health) was injected intraperitoneally at 48-hour intervals to induce superovulation. Mice were euthanized by cervical dislocation 15-17 hours after hCG injection, and cumulus-oocyte complexes were collected in the KSOM^AA^BSA medium (MR-106-D; Merck, Darmstadt, Germany) equilibrated with 6% CO_2_ in air. Then, oocytes were isolated from the cumulus-oocyte complex by treating with 3 µg/ml hyaluronidase (H3506; Sigma-Aldrich, St Louis, MO, USA) for approximately 5 min.

Strontium activation of oocytes was performed by treating oocytes for 30–40 min with a strontium solution containing 5 mM SrCl_2_ (193-09442; Wako Pure Chemical Industries, Osaka, Japan) and 2.25 mM EGTA (E8145; Sigma-Aldrich, St Louis, MO, USA) dissolved in the KSOM^AA^BSA medium as previously described (Kishigami & Wakayama, 2007).

### Preparation of mRNA probes

mRNA probes for histone H2B-mCherry, RCC1-EGFP, and Nup153-EFGP were generated as previously (Suzuki et al., 2019), and that for superfolder GFP-nucleoplasmin (NPM)-NLS (sfGFP-NLS) was generated as previously described (Oda et al., 2023). Briefly, cDNA encoding a protein of interest fused with fluorescent protein was cloned into pcDNA3.1 containing a poly(A) tail sequence for mRNA synthesis. Cloned plasmids were linearized at the *Xho* I sites. mRNAs were synthesized using RiboMAX^TM^ Large Scale RNA Production Systems-T7 (Promega, Madison, WI, USA) using the corresponding plasmid as a template as previously described (Yamagata et al., 2005). For efficient translation of the fusion proteins in oocytes, the 5′-end of each mRNA was capped using Ribo m^7^G Cap Analog (Promega), according to the manufacturer’s protocol. To circumvent integration of template DNA into the genome, the reaction mixtures for *in vitro* transcription were treated with RQ-1 RNase-free DNase I (Promega). Synthesized mRNAs were treated with phenol–chloroform to remove protein components. The mRNAs were further purified by filtration using MicroSpin^TM^ S-200 HR columns (Amersham Biosciences, Piscataway, NJ, USA) to remove unreacted substrates (RNA reaction intermediates) and then stored at −80°C until use.

### Injection of mRNA probes

Microinjection of mRNA probes into oocytes was performed as described previously (Yamagata, 2010). Briefly, mRNAs were diluted to 5 ng/μl each for histone H2B-mCherry and RCC1-EGFP, 100 ng/µl for Nup153-EGFP and 30 ng/μl for sfGFP-NPM-NLS-L using ultrapure water (Thermo Fisher Scientific Barnstead Smart2Pure), and an aliquot (0.3 μl) was placed in a micromanipulation chamber. Oocytes were transferred to 8 μL drops of HEPES-buffered Chatot-Ziomek-Bavister (HEPES-CZB) medium (Chatot et al., 1989). The mRNA solution was aspirated into a narrow glass pipette (1 μm diameter) after washing with HEPES-CZB contains 12% Polyvinylpyrrolidone (PVP-HEPES-CZB) medium and injected into the oocyte using a piezo-driven manipulator. The mRNA-injected oocytes were incubated at 37°C under 6% CO_2_ in air for at least 3 h for the translation.

### Injection of DNA

DNA was purified by phenol/chloroform treatment, and dissolved in ultrapure water to final concentrations of 20, 100, 250, and 500 ng/µl. The DNA solutions were aliquoted in 0.3 µl and stored at −30°C until use. For microinjection of DNA, a DNA aliquot (0.3 μl) was placed in a micromanipulation chamber. Oocytes were transferred to the 8 µl drop of HEPES-CZB medium in the chamber. After the washing the injection pipette (1 μm diameter) with HEPES-CZB medium containing 12% PVP, the DNA solution was aspirated into the pipette and injected into the oocyte using a piezo-driven manipulator. A few picoliters of solution (equivalent to a sphere with a radius of approximately 10 µm) were introduced into the cytoplasm, and the pipette was removed gently. The mRNA-injected oocytes were incubated at 37°C under 5% CO_2_ in air for at least 3 h to wait for protein production.

### Live-cell imaging

Oocytes were transferred to 5 μl drops of KSOM^AA^ medium containing 0.00025% Polyvinyl alcohol (PVA) and 100 μM EDTA on a glass-bottomed dish (P35G-1.5-14-C; Mat Tek Corp., Ashland, MA, USA). The oocytes were imaged using a spinning-disk confocal fluorescence microscopy system (CSU-W1; Yokogawa Electric Corp., Tokyo, Japan) equipped with a temperature-controlled chamber (STXG-IX3WX-SET; Tokai Hit Inc., Shizuoka, Japan) set at 37°C in 6% CO_2_ and 5% O_2_ with saturated humidity. The inverted microscopy (IX-73, Evident Scientific, Inc., Tokyo, Japan) was equipped with an EM-CCD camera (iXon3 DU897E-CS0-#BV-Y; Andor Technology Ltd., Belfast, UK), focus control devise (Mac6000; Ludl Electronic Products Ltd., NY, US) and x-y stage control devise (SIGMAKOKI CO., LTD., Tokyo, Japan). Fluorescence images were taken with a 40x objective lens (UPLSAPO40XS, NA = 1.25) and laser lines at 488 nm (0.05 mW) and 555 nm (0.1 mW) (LDI-PRIME LASER DIODE ILLUMINATOR; 89North, VT, USA) at 105 ms exposure time. Images were taken for 2 days at 10- or 15-minute intervals. For each time point, 46 different focal plane images were taken at 2 µm intervals. The system was controlled by µManager 2.0 software (Edelstein et al., 2014). The visualization and image analyses were performed using ImageJ/Fiji image analysis platform (https://imagej.net/Fiji; Schindelin et al., 2012).

### Immunofluorescence staining

Oocytes were fixed with 4% paraformaldehyde (163-20145; Wako Pure Chemical Industries, Osaka, Japan) for 30 min at room temperature (RT; approximately 26°C). After fixation, the oocytes were washed twice with phosphate-buffered saline (PBS) containing 0.02% PVA. The oocytes were permeabilized with 0.25% Triton X-100 in PBS for 20 min at RT, washed twice with blocking solution (PBS containing 3% bovine serum albumin (BSA) and stored for 60 min in the blocking solution at RT. The oocytes were then incubated with primary antibodies dissolved in the blocking solution at 4°C for more than 8 h; the primary antibodies used are anti-histone H2A (1:100 dilution), anti-histone H3 (1:400 dilution), anti-histone H4 (1:100 dilution), anti-NPC (1:500 dilution) and anti-Ran (1:100 dilution). After being washed twice in same solution, the oocytes were further incubated with secondary antibodies (1:500 dilution) for 60 min at RT. After the washing with PBS, the embryos were transferred to PBS containing 0.5 µg/ml of 4′,6-diamidino-2-phenylindole (DAPI, 19178-91; Nacalai Tesque Inc., Kyoto, Japan) in glass bottomed-dish (P35G-1.5-14-C; Mat Tek Corp., Ashland, MA, USA). Immunostaining with anti-B23 antibody was performed according to the method of (Shishova et al., 2015). Briefly, oocytes were fixed in 4% PFA for 30 min at RT, washed twice with PBS containing 0.02% PVA, permeabilized with 0.5% Triton X-100 in PBS, and treated with 1 µg/ml proteinase-K (29442-14; Nacalai Tesque Inc., Kyoto, Japan)/PBS containing 0.02% PVA for 60 min at RT. After blocking overnight with blocking solution, the oocytes were treated by primary and secondary antibodies with same procedures described above. The oocytes were observed using a super-resolution spinning-disk confocal scanner microscope system (CSU-W1 SoRa, Yokogawa Electric Corp., Tokyo, Japan) equipped with a sCMOS camera (Pime95B; Teledyne Photometrics, Tuscon, AZ, USA) as previously described (Hatano et al., 2022). Images were taken at 101 different focal planes at 0.5 µm intervals with 405nm, 488nm and 561nm laser lines (OBIS; Coherent, CA, USA) at 105 ms exposure time, using a 40x objective (UPLSAPO40XS; NA=1.25, OLYMPUS, Tokyo, Japan).

### Electron microscopy

Oocytes were fixed overnight at 4°C with 2.5% (w/v) glutaraldehyde (3043; TAAB Laboratory Equipment, Ltd., Reading, UK) and washed twice in 0.1 M PB (pH 7.4) containing 0.1 % (w/v) PVA. The fixed oocytes were embedded in 50 μl droplets of 0.5% agarose (SeaPlaque™ Agarose, Lonza, Basel, Switzerland; 50101) in 0.1M PB on a glass-bottomed microwell dish (P35G-1.5-10-C; Mat Tek Corp., Ashland, MA, USA) and centrifuged at 2000 rpm for 2 min. The embryos were post-fixed with 1.5% OsO_4_ (157-00404; Fujifilm Wako, Tokyo, Japan), stained with 2% (w/v) uranyl acetate (8473-1M; Wako Pure Chemical Industries) for 1 h, dehydrated and embedded in Epok812 (02-1001; Okenshoji Co. Ltd., Tokyo, Japan). Ultra-thin sections (80 nm thick) were prepared using an ultramicrotome (UC7, Leica Microsystems, Wetzlar, Germany) and stained with 4% uranyl acetate, followed by a commercial ready-to-use solution of lead citrate (18-0875-2; Sigma-Aldrich). Electron microscope (EM) images were acquired using a JEM-1400plus electron microscope (100 kV; JEOL, Tokyo, Japan).

### Statistical analysis

The statistical analysis was performed using R version 4.1.3 (R Core Team, 2017) and the stats package. *p*-values were calculated using the Steel-Dwass test for comparing independent proportions. *p*<0.05 was considered statistically significant.

## Acknowledgments

We thank Drs. Noriko Saitoh and Kenji Watanabe for valuable discussions on the structure of the nucleolus. This study was supported by JSPS Kakenhi Grant Number JP21K15103 to HO, JP18H05526, JP21H04764, and JP24H02325 to HK, JP18H05533, JP20H00454, and JP23K05636 to YH, JP18H05528 to TH, and JP18H05528 and JP24H02325 to KY.

## Author contributions

NY, HK, YH, TH, and KY designed the experiments. NY, TS, and HO performed the experiments. All authors analyzed and discussed the data, and NY, YH, TH, and KY wrote the manuscript.

