## Supplementary Figures for "Reconstruction of artificial nuclei with nuclear import activity in living mouse oocytes"

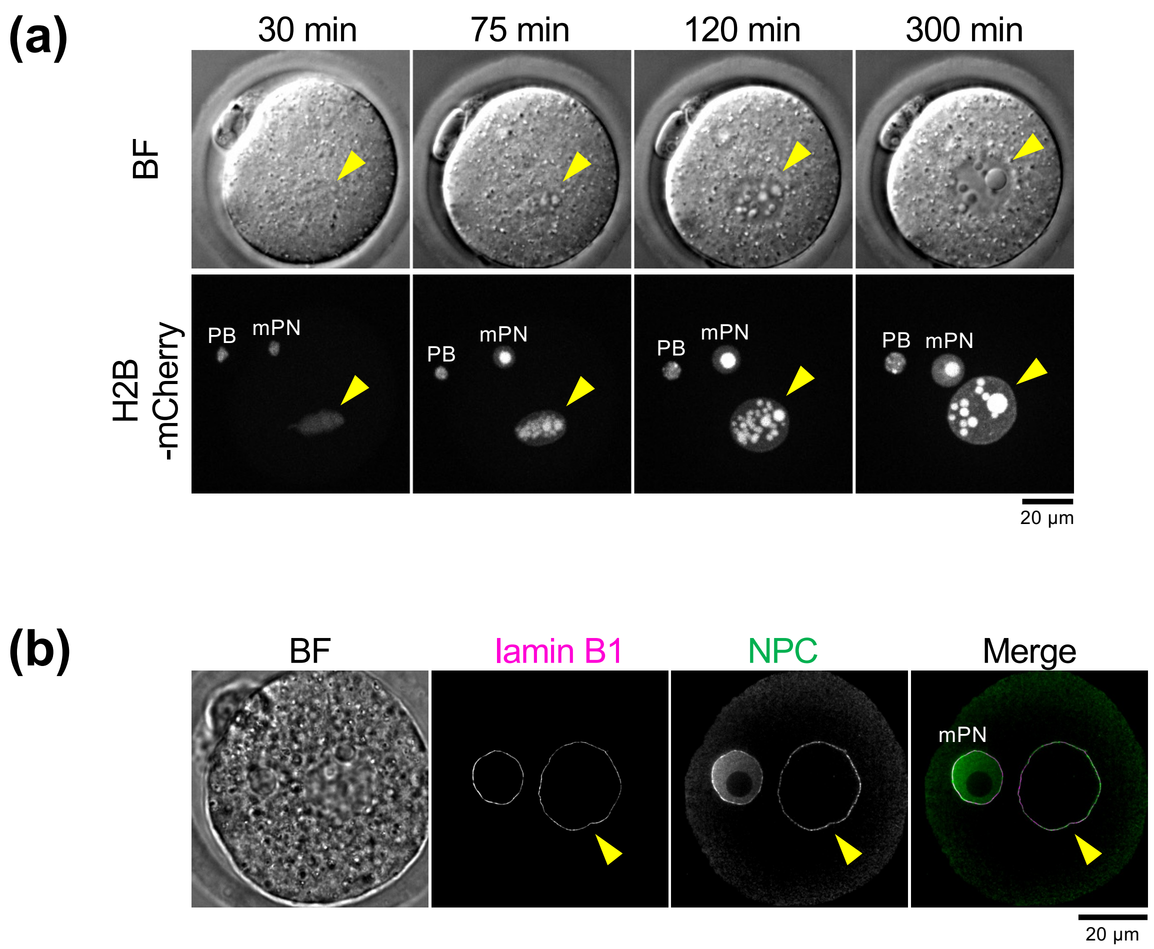


**Figure S1. Injection of λDNA forms nuclear-like structures in mouse oocyte.**
(a) Time-lapse fluorescence images of living mouse oocytes. Anaphase II oocytes expressing H2B-mCherry were injected with 250 ng/µl λDNA. Bright-field (BF) (upper panels) and H2B-mCherry (lower panels) images are shown. The numbers above each image indicate the time in minutes after DNA injection. Single focus images are shown for BF and maximum projection images for H2B-mCherry. Yellow arrowheads indicate the region of injected λDNA. PB, polar body. mPN, maternal pronucleus. Scale bars, 20 µm.
(b) Immunostaining images of mouse oocytes using anti-lamin B1 and anti-nuclear pore complex (NPC) mAb414 antibodies, respectively. Images were taken 300 min after injecting 250 ng/µl λDNA into anaphase II oocytes. Images are single planes of the nuclei. The leftmost image is a BF image. The merged image, lamin B1 (magenta) and NPC (green). Yellow arrowheads indicate the region of injected λDNA. mPN, maternal pronucleus. Scale bar, 20 µm.

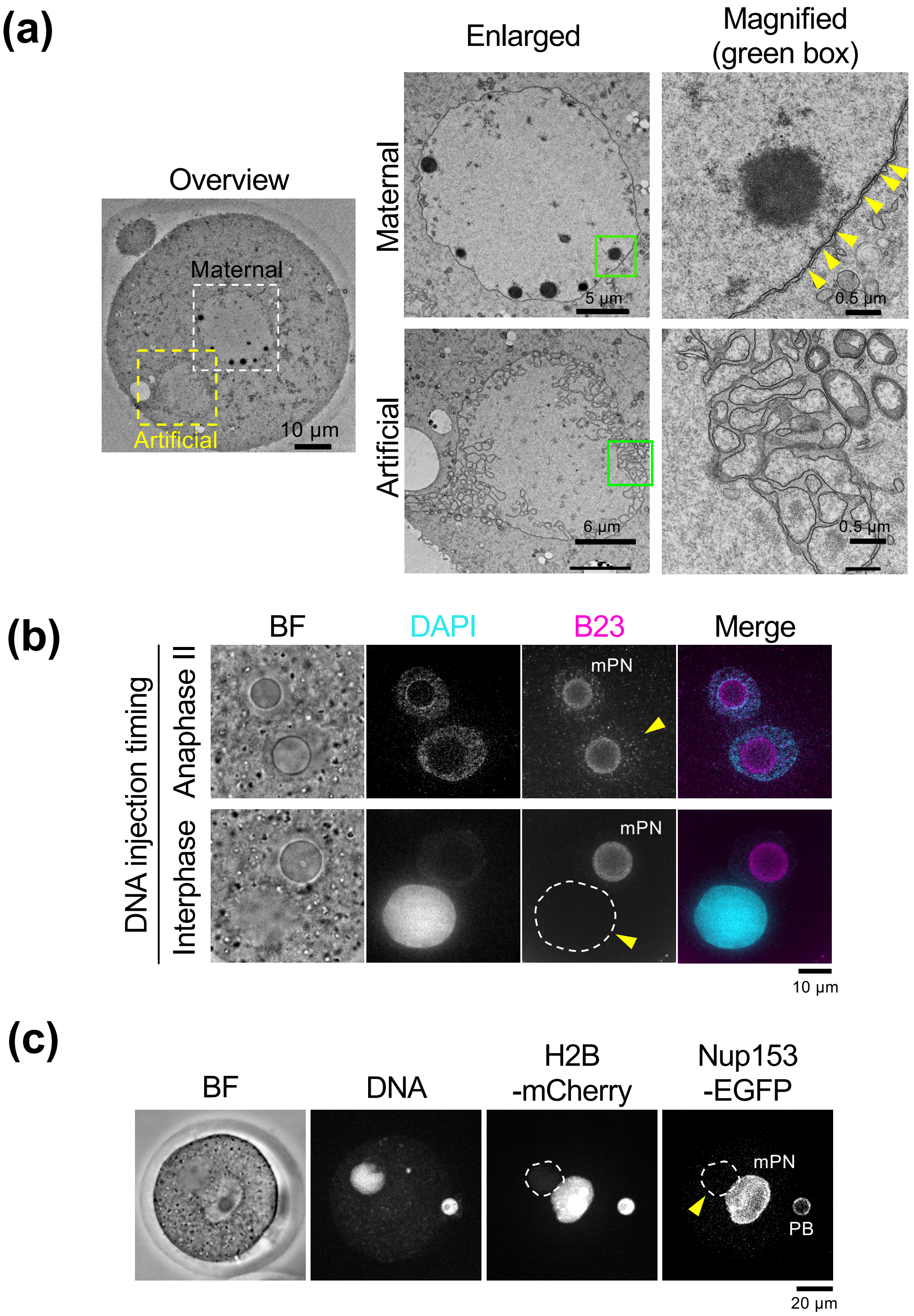


**Figure S2.**

(a) Electron microscopy (EM) images of mouse oocytes 300 min after injecting 250 ng/µl T4 DNA at interphase. The leftmost image is an overview EM image. White- and yellow-boxed regions in the overview are enlarged in the second left panels ("Enlarged" panels). Upper and lower images are the regions of maternal pronuclei (maternal) and injected T4 DNA (artificial), respectively. The rightmost panels are magnified images of the green-boxed regions on the left. Yellow arrowheads indicate the nuclear pore complex (NPC)-like structures on the nuclear envelope. Scale bars are indicated in each panel.

(b) Immunostaining images of mouse oocytes using anti-B23 antibody. Images were taken 300 min after injecting 250 ng/µl T4 DNA into anaphase II (upper panels) and interphase (lower panels) oocytes. DNA was stained with DAPI. The images are single planes of the nuclei. The leftmost image is a BF image. The merged image, DNA (cyan) and B23 (magenta). mPN, maternal pronucleus. Yellow arrowheads indicate the regions of DNA injection. The dashed area indicates the region corresponding to the signal of DAPI. Scale bar, 10 µm.
(c) Images of living mouse oocytes. Interphase oocytes expressing Nup153-EGFP and H2B-mCherry were injected with 250 ng/µl T4 DNA. oocytes were fixed at 300 min after injecting T4 DNA. DNA was stained with Hoechst 33342. A single plane image is shown for BF and maximum projection images for DNA, H2B-mCherry, and Nup153-EGFP. The dashed area is the region where DNA was injected. PB, polar body. mPN, maternal pronucleus. Scale bar, 20 µm.
